# CDP-DAG synthases regulate plant growth and broad-spectrum disease resistance

**DOI:** 10.1101/2025.01.24.634677

**Authors:** Ronglei Tan, Gan Sha, Qiuwen Gong, Lei Yang, Wei Yang, Xiaofan Liu, Yufei Li, Jiasen Cheng, Xin Qiao Du, Hongwei Xue, Qiang Li, Jie Luo, Guotian Li

**Author notes:** These authors contributed equally to this article (Ronglei Tan, Gan Sha). Correspondence: Guotian Li.

## Abstract

Phosphatidic acid (PA) functions as a cell membrane component and signaling molecule in plants. PA metabolism has multiple routes, in one of which PA is converted into cytidine diphosphate diacylglycerol (CDP-DAG) by CDP-DAG synthases (CDSs). *CDS* genes are highly conserved in plants. Here, we found that knock-down of the *CDS* gene enhanced the resistance of *Arabidopsis thaliana* to multiple pathogens, with a growth penalty. When *Arabidopsis* leaves were treated with chitin or flg22, reactive oxygen species (ROS) production in *cds* mutants was significantly higher than that in the wild-type (WT). Similarly, phosphorylation of mitogen-activated protein kinases (MAPKs) in the *cds1cds2* double mutant was significantly increased compared to the WT. By integrating lipidomics, transcriptomics, and metabolomics data, PA accumulation was observed in mutants *cds1cds2,* activating the jasmonic acid (JA) and salicylic acid (SA) signaling pathway, and increasing transcript levels of plant defense-related genes. Significant accumulation of the downstream metabolites including serotonin and 5-methoxyindole was also found, which plays important roles in plant immunity. In conclusion, our study indicated the role of CDSs in broad-spectrum disease resistance in *Arabidopsis* and that CDSs are involved in plant metabolic regulation.

## 1. INTRODUCTION

Phospholipids, as amphipathic molecules, are major structural components of biological membranes and play vital roles in membrane trafficking. Some phospholipids such as phosphatidic acid (PA), function in signal transduction (Xing et al., 2021). Phospholipids are involved in plant growth, development, and responses to biotic stress, by binding downstream proteins through lipid-protein interactions and modulating their expression levels, subcellular localization, or protein functions (Ali et al., 2022).

PA, the simplest yet key phospholipid molecule, is highly dynamic and relays multiple cellular signaling events, including responses to biotic stress (Yao et al., 2024). In plants, PA interacts with proteins that function in or outside cell nucleus to regulate cellular and physiological processes and plays indispensable roles in plant immunity (Xing et al., 2021). PA binds to respiratory burst oxidase homolog D (RBOHD), which generates reactive oxygen species (ROS) in plant immunity, including both the pattern-triggered immunity (PTI) and effector-triggered immunity (ETI) processes. The binding of PA to RBOHD stabilizes RBOHD, thereby modulating ROS production (Qi et al., 2024, Zhang et al., 2009). Diacylglycerol kinase 5 (DGK5) catalyzes the synthesis of PA from diacylglycerol, and recent studies show that pathogen infection activates DGK5 to synthesize PA, which inhibits RBOHD protein degradation, thereby promoting ROS production and enhancing plant immunity (Kong et al., 2024, Qi et al., 2024). In addition, PA binds to and inhibits abscisic acid (ABA) biosynthesis enzyme ABA2 and suppresses ABA production in response to abiotic stress in *Arabidopsis* (Li et al., 2024). It is clear from these studies that PA plays an important role in stress responses in *Arabidopsis*.

As a dynamic molecule, PA is synthesized through multiple biochemical pathways, including the *de novo*-Kennedy pathway, and pathways involved in DGKs and phospholipase D (PLD) (Li and Wang, 2019). PA is the central precursor for all glycerophospholipids and can be converted to cytidine diphosphate diacylglycerol (CDP-DAG), a vital metabolic intermediate, through the CDP-DAG synthases (CDSs), and then towards PI, phosphatidylglycerol (PG) and cardiolipin (CL) via different enzymatic reactions (Jennings and Epand, 2020). There are five *CDS* genes in *Arabidopsis*, complete loss of expression of both *CDS1* and *CDS2* cause seedling lethality. *CDS3* is specifically expressed in certain plant structures such as stamens and mature pollen (Zhou et al., 2013). CDS4 and CDS5 are localized at the plasma membrane, and are important for PG synthesis in chloroplast-like vesicle structures (Haselier et al., 2010). To study *Arabidopsis CDS1* and *CDS2* in detail, the *cds1cds2* knock-down line was generated (Du et al., 2022). This line exhibit reduced auxin levels and aberrant polar distribution of PIN1, leading to delayed embryonic development. Furthermore, the *cds1cds2* mutant show enhanced resistance to the oomycete pathogen *Phytophthora capsici* (Sha et al., 2023). The *CDS1* homologue from rice, *RBL1*, negatively regulates rice immunity. When *RBL1* is deleted or mutated in rice, a strong autoimmune response is observed, with an increased level of PA and decreased levels of PI and its derivatives. Rice *rbl1* mutants show enhanced resistance to the rice blast fungus *Magnaporthe oryzae* and the bacterial blight pathogen *Xanthomonas oryzae* pv. *oryzae* (Sha et al., 2023). However, how CDSs regulate plant growth, development and immunity is largely unknown. Here, we investigated the role of *Arabidopsis* CDSs in plant growth and broad-spectrum disease resistance by integrating transcriptomics, lipidomics and metabolomics approaches.

## 2. MATERIALS AND METHODS

### 2.1 Plant materials, strains and growth conditions

The *Arabidopsis thaliana* ecotype Columbia (Col-0) was used as the background of all lines used in this study. The *cds1* (SALK_001496) and *cds2* (SALK_106246) were obtained from Nottingham Arabidopsis Stock Centre. The *cds1cds2* double mutant was obtained by crossing homozygous *cds1* and *cds2.* All seedlings were grown in potting soil mix (peat soil: rich soil: vermiculite of 1:1:1, v/v/v) in a humidity plant growth chamber at 22 °C, 60% humidity, with a photoperiod of 16:8 hours (h) light: dark (Du et al., 2022). *Botrytis cinerea* B05.10 (*B. cinerea*) and *Sclerotinia sclerotiorum* 1980 (*S. sclerotiorum*) were obtained from Prof. Long Yang (Kamaruzzaman et al., 2019) and Prof. Jiasen Cheng (Gong et al., 2022), respectively. The *S. sclerotiorum* strain was cultured on potato dextrose agar (PDA) plates at 25 °C for 3-4 d in the dark. The *B. cinerea* strain was cultured on PDA plates at 20 °C for 2-3 d in the dark. *Pseudomonas syringae* pv. *tomato* DC3000 (*Pst* DC3000) was obtained from Prof. Xiufang Xin (Yuan et al., 2021). *Pst* DC3000 strains were incubated with 5 mL liquid LB medium at 28 °C overnight, reaching a cell density of OD_600_ = 0.8∼1.0.

### 2.2 Plant infection assays

Infection assays of *Arabidopsis* with *S. sclerotiorum* and *B. cinerea* were conducted as previously described with minor modifications (Gong et al., 2022, Kamaruzzaman et al., 2019). Four-week-old leaves of *Arabidopsis* were used for pathogen inoculation. Fully extended *Arabidopsis* leaves were detached, surface-sterilized with 75% ethanol and placed on 0.8% water agar plates. Subsequently, mycelial blocks (2 mm × 2 mm) of *S. sclerotiorum* and *B. cinerea* were inoculated in the center of each leaf. The infected leaves were then kept at 25 °C in the dark. At 30 hours post-inoculation, the infected leaves were photographed and the lesion areas were measured. Infection assays with *Pst* DC3000 were conducted as previously described (Hu et al., 2022). *Pst* DC3000 were incubated with 5 mL liquid LB medium at 28 °C overnight, reaching a cell density of OD_600_ = 0.8∼1.0. Bacteria cultures were then harvested through centrifugation (4000 rpm, 25 °C for 5 min), washed with sterile water, and diluted to a cell density of OD_600_ = 0.0005. The bacteria suspensions were injected into the leaves using a needleless syringe, and the plants were subsequently covered with a clear plastic cover to maintain high humidity levels for 6 h. For each biological replicate two leaf discs (6 mm in diameter) were obtained from different leaves and surface-sterilized with 75% ethanol. The leaf discs were mashed and suspended in sterile water, the bacterial suspensions of different dilution ratios were prepared and then plated on TSA agar plates supplemented with 50 mg/L rifampicin. Colonies were counted after 48 h of incubation at 28 °C.

### 2.3 Measurement of ROS

ROS was measured as previously described (Liu et al., 2017). For ROS burst assay, 3-week-old leaves of the *Arabidopsis* plants were excised into 3 mm × 3 mm discs. These leaf discs were submerged in double-distilled sterile water in a 96-well plate and incubated overnight under controlled light conditions to counteract the wounding effects. After the incubation was finished, distilled water was replaced with a solution containing 50 μΜ luminol, 10 μg/ml horseradish peroxidase, and indicated elicitors (either 120 nM chitin or 200 nM flg22) or distilled water (as the blank). Chemiluminescence was measured at 500-ms intervals continuously over a period of 40 min on a SPARK-10M microplate reader (TECAN).

### 2.4 RNA extraction and RT–qPCR assays

Leaves from 3-week-old *Arabidopsis* plants for each genotype were excised and immediately snap-frozen in liquid nitrogen. Total plant RNA was isolated using TRIzol (Vazyme). The separation of the organic and aqueous phases is facilitated with chloroform. The total RNA was precipitated from the aqueous phase using Isopropanol. HiScript II Q RT SuperMix for qPCR (Vazyme) was used to remove residual genomic DNA and synthesize complementary DNA (Sha et al., 2023). The RT-qPCR analysis was performed using SYBR Green mix with a Bio-Rad CFX96 Real-Time System coupled to a C1000 Thermal Cycler. All the primer information of RT–qPCR assays were provided in Supplementary Table 1. *Arabidopsis MPK3* (AT3G45640) and *MPK4* (AT4G01370) genes encode two mitogen-activated protein kinases (MAPK), play critical roles in plant disease resistance by regulating multiple defense responses. *Arabidopsis ACTIN2* (AT3G18780) gene was used as the internal control to normalize the expression level of *MPK3* and *MPK4* genes for gene expression analysis. Furthermore, the expression level of target genes was quantitatively assessed using the 2 ^-ΔΔ*C*^*^T^* method (Livak and Schmittgen, 2001).

### 2.5 MAPK phosphorylation assays

Two-week-old *Arabidopsis* plant grown on 1/2 MS plates were used for MAPK phosphorylation analysis. We placed 4-6 seedlings into sterile water overnight before the treatment, then immersed the roots of the seedlings in a solution containing indicated elicitors (120 nM chitin or 200 nM flg22) and incubated for 15 min. Each sample was collected in a centrifuge tube with liquid nitrogen and thoroughly ground to fine powder. The samples were solubilized in the extraction buffer (50 mM Tris-HCl, pH 7.5, 150 mM NaCl, 1 mM EDTA, 1% Triton X-100, 0.5 mM PMSF and 1% protease inhibitor cocktail) and boiled at 95 °C for 10 min. The protein extracts were centrifuged at 12,000 rpm at 4 °C for 5 min (Zhai et al., 2022). A 20-μl mixed sample (containing 14-μl protein extract, 2 μl 0.4 M DTT, 4 μl 5×SDS loading buffer) was loaded on a 10% SDS–PAGE gel. The fractioned proteins were transferred to the PVDF membrane for immunoblotting. Blots were blocked with 5% BSA prepared in TBST at room temperature for 1 h, and then incubated with anti-phospho-p44/42 antibody (Cell Signaling Technology, 1:2000 dilution) overnight at 4 °C. Subsequently, the blots were probed with a secondary antibody Goat anti-rabbit IgG HRP (Cell Signaling Technology, 1:5000 dilution) at room temperature for 1 h. The phosphorylated MAPK (pMAPKs) proteins were then visualized using ECL Chemiluminescence Kit (Vazyme., catalogue no. E423) on the ChemiDoc Touch Imaging System (Bio-Rad). *Arabidopsis* ribulose bisphosphate carboxylase/oxygenase (RUBISCO) was used as the internal control (Yuan et al., 2021).

### 2.6 RNA-seq analysis

Leaf samples from 4-week-old *Arabidopsis* plants were used for RNA-seq analysis. The 2×150-bp paired-end sequencing was performed on the BGISEQ-500 sequencer (BGI, Shenzhen, China). Raw sequencing data was filtered by removing sequencing adapters, low-quality reads (base quality less than or equal to five) and reads with high percentage of unknown bases (‘N’ base > 5%), and the resultant clean data were stored in the FASTQ format for downstream analysis. Clean reads were aligned against the *Arabidopsis* Col-0 genome using HISAT2 (Huang et al., 2017). Quantification of gene expression was performed using FeatureCounts. DESeq2 (version: 1.32.0) was used to assess differential expression between sample groups. Differentially expressed genes (DEGs) were identified by applying a |Fold Change|>1 and false discovery rate (FDR) < 0.05. DEGs were functionally annotated using the GO database (http://geneontology.org/) (Yu et al., 2012).

### 2.7 Lipid extraction and analysis

Leaf samples from 4-week-old *Arabidopsis* plants were used for phospholipid extraction. Briefly, around 10 mg of fresh *Arabidopsis* leaf tissues were incubated in preheated isopropanol (75 °C) containing 0.01% butylated hydroxytoluene (BHT) for 15 min. After cooling to room temperature, chloroform was added to the samples, and then incubated with shaking at room temperature for 1 h. Lipid extracts were transferred to new glass tubes to repeat the extraction procedure until the leaf sample became bleached. 1M KCl and water immediately added, and the upper phase was discarded after centrifugation (4 °C, 1,000g for 10 min). The lipid extracts were combined and evaporated under nitrogen gas, and then redissolved in chloroform to a final concentration at 10 mg/ml (Hong et al., 2018, Zhao et al., 2024). Phospholipids were measured with an electrospray ionization source in positive-ion mode (Lu et al., 2019).

### 2.8 Metabolite profiling analysis

Freeze-dried tissue samples were ground into fine powder in a ball mill, and 50 mg of powder for each sample was mixed with 5 mL methanol/water (3:1, v/v), and incubated at 4 °C overnight for water-soluble metabolites. Then, all samples were centrifuged at 10,000 g at 4 °C for 10 min. The extracts were absorbed (CNWBOND Carbon-GCBSPEC cartridge) and filtered before liquid chromatography tandem-mass spectrometry (LC-MS/MS) analysis (Zhang et al., 2020). The extracted samples were assayed by a Quadrupole Time of Flight (QTOF) system (TripleTOF 5600+; AB Sciex, USA) for untargeted detection and a QTrap system (QTRAP 6500+; AB Sciex) for targeted detection as described previously before (Zhou et al., 2022).

### 2.9 Quantification of total salicylic acid (SA) in leaves

Leaf samples from 4-week-old *Arabidopsis* plants were used for total SA analysis. Leaf tissues were collected and subjected to freeze-drying to preserve their biochemical content. Approximately 20 mg of the freeze-dried leaf tissues were pulverized using liquid nitrogen, then mixed with 1 ml of 80% methanol and sonicated at 70 °C for 15 minutes to extract the SA. After centrifuging the mixture at 7,000g for 4 °C, the supernatant was collected and passed through a filter to remove any remaining particulates. The filtrate, along with serial dilutions of the SA standard, was analyzed using high-performance liquid chromatography (HPLC) coupled with tandem mass spectrometry (MS/MS). The concentration of SA in each sample was calculated by comparing the results to a calibration curve generated from the SA standard (Sha et al., 2023).

## 3. RESULTS

To investigate the function of *CDS1* and *CDS2* in growth regulation and immunity of *Arabidopsis*, the growth phenotype of *Arabidopsis* mutants with reduced transcription levels of *CDS1* and *CDS2* caused by T-DNA insertions (Du et al., 2022) at mature stage was observed. At 40 days post-sowing, the height of the *cds1cds2* mutant (26.04 cm) was significantly lower than that of the WT (35.48 cm) (Fig. 1a), with a 27% reduction (Fig. 1b). The number of siliques in *cds1cds2* was significantly reduced by 51%, compared to that in the WT. The plant height and the number of siliques per plant in the *cds1* and *cds2* single mutant lines showed no difference relative to the WT (Fig. 1c, d).

**FIGURE 1.**
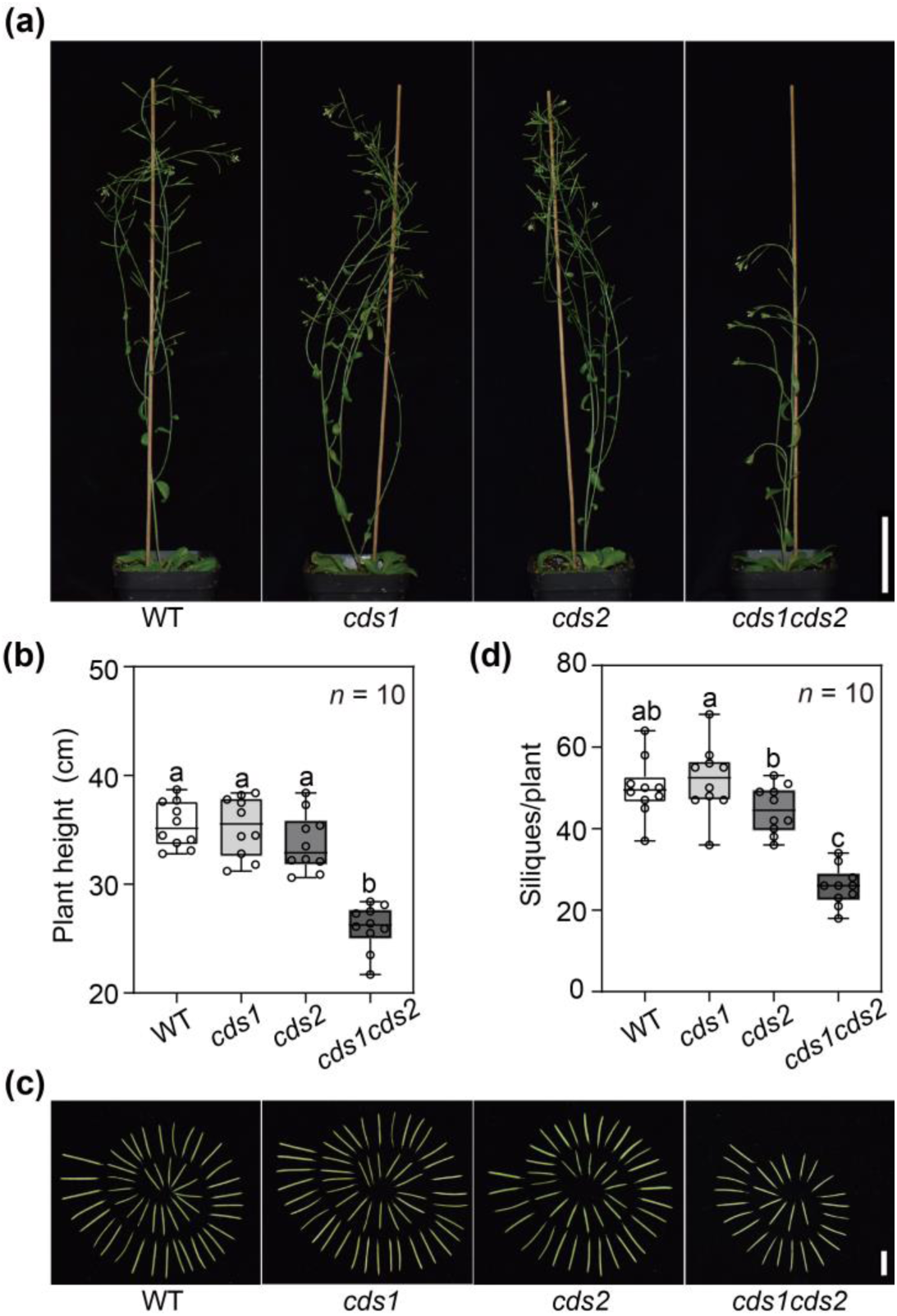
Plant growth is reduced in *Arabidopsis cds1cds2* mutants. (a) The *cds* mutant and wild-type (WT, Col-0) plants at 40 days post-sowing (dps). Bar, 5 cm. Plants were grown in a growth chamber at 22 °C, 60% humidity with a photoperiod of 16:8 hours (h) light: dark. (b) Plant height of *cds* mutants and WT. (c) Photograph of siliques from a representative plant and (d) the number of siliques per plant at 40 dps. Bar, 1 cm. Data are displayed as box and whisker plots with individual data points. The box plot elements are: center line, median; box limits, 25^th^ and 75^th^ percentiles. *n* indicates the number of biological replicates. Letters indicate statistically significant differences (ANOVA, *p*< 0.05). Duncan’s Multiple Range Test as method of post-hoc test.

The infection assays of 4-week-old *cds* mutant lines aimed to further investigate the resistance levels of these mutants to multiple pathogens. In the assay, the mean lesion area caused by *S. sclerotium* in the *cds1cds2* mutant was 0.22 mm^2^ (Fig. 2a), which was 47% smaller than that of the WT (0.41 mm^2^) (Fig. 2a). In contrast, enhanced resistance to *S. sclerotium* has not been observed in single *cds* mutants, which mean lesion area were 0.41 mm^2^ and 0.34 mm^2^, respectively. The *cds1*, *cds2*, and *cds1cds2* mutant plants all exhibited enhanced resistance to *B. cinerea* (Fig. 2b). The *cds1cds2* mutants, the *cds1* mutants, and the *cds2* mutants showed 32%, 27%, and 28% reductions, respectively, in lesion area compared to the WT (Fig. 2b). *Pst* DC3000 infection assay showed that the *cds1cds2* mutants exhibited slightly smaller brown leaf spots compared to the WT and other lines (Fig. 2c). Additionally, the colony-forming units (CFU) assay showed that the multiplication of *Pst* DC3000 was greatly restricted in *cds1cds2* mutants. The bacterial density within the leaves of *cds1cds2* mutants was 1.8 x 10^5^ CFU/cm^2^, decreased by 82.3%, 82.2%, and 83.4% compared to that of WT (1.02 x 10^6^ CFU/cm^2^), *cds1* mutant (1.01 x 10^6^ CFU/cm^2^), and *cds2* mutant (1.08 x 10^6^ CFU/cm^2^), respectively (Fig. 2c). In summary, the *cds1cds2* mutant lines showed resistance to multiple pathogens.

**FIGURE 2.**
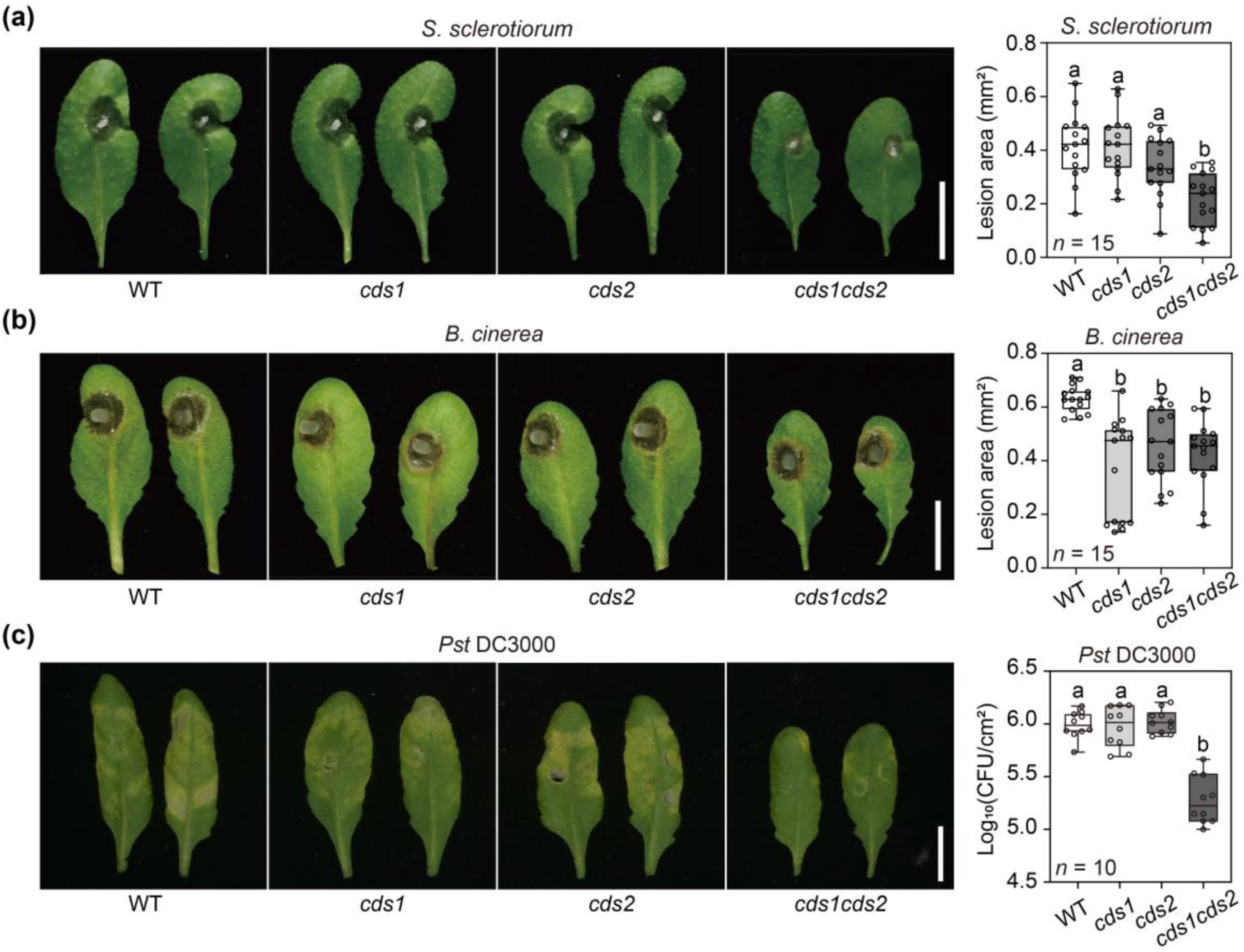
*Arabidopsis cds1cds2* mutants show broad-spectrum pathogen resistance. (a) Infection assays with the *Sclerotinia sclerotiorum* (*S. sclerotiorum*) and lesion areas of *cds* mutant lines at 30 hours post-inoculation (hpi); bar, 1 cm. (b) Infection assays with the *Botrytis cinerea* (*B. cinerea*) and lesion areas at 30 hpi. (e) Bacterial infection assays with *Pseudomonas syringae* pv. *tomato* DC3000 (*Pst* DC3000) at 3 days post-inoculation (dpi) and related bacterial populations. Box plot elements are: center line, median; box limits, 25^th^ and 75^th^ percentiles. *n* indicates the number of biological replicates. Letters indicate statistically significant differences (ANOVA, *p*< 0.05). Duncan’s Multiple Range Test as method of post-hoc test.

The ROS bioassay of *cds* mutant lines, treated with pathogen-associated molecular patterns (PAMPs), was conducted to investigate the potential mechanisms underlying the enhanced disease resistance of the *cds1cds2* mutant. Under treatments of chitin and flg22, the ROS level in the *cds1cds2* mutant was 2.64- and 1.31-fold higher than that of the WT, respectively (Fig. 3a). The total number of photons in the *cds1cds2* mutant showed 2.72- and 1.33-fold increase compared to that in the WT (Fig. 3b). PTI often activates the downstream MAPK cascade immune response. The expression levels of *MPK3* and *MPK4* genes in *cds* mutant leaves were significantly upregulated compared to the WT (Fig. 3c). Furthermore, total proteins from *cds* mutant leaf tissues treated with PAMPs were extracted to analyze the levels of pMAPKs using immunoblotting (Kong et al., 2024). The accumulation levels of pMAPKs, the *cds1cds2* mutant showed significantly higher levels of pMAPKs than the WT (Fig. 3d). SA, a key signaling molecule in plant immunity, induces disease resistance to various viruses, fungi and bacteria in plants. A significant increase in SA levels was detected in *cds1cds2* mutants, implying the important role of SA in enhancing disease resistance in *cds1cds2* mutants (Fig. S1). These results indicated that the *cds1cds2* mutant displayed a more robust immune response than the WT, and the enhanced disease resistance was associated with increased levels of ROS, pMAPKs and SA.

**FIGURE 3.**
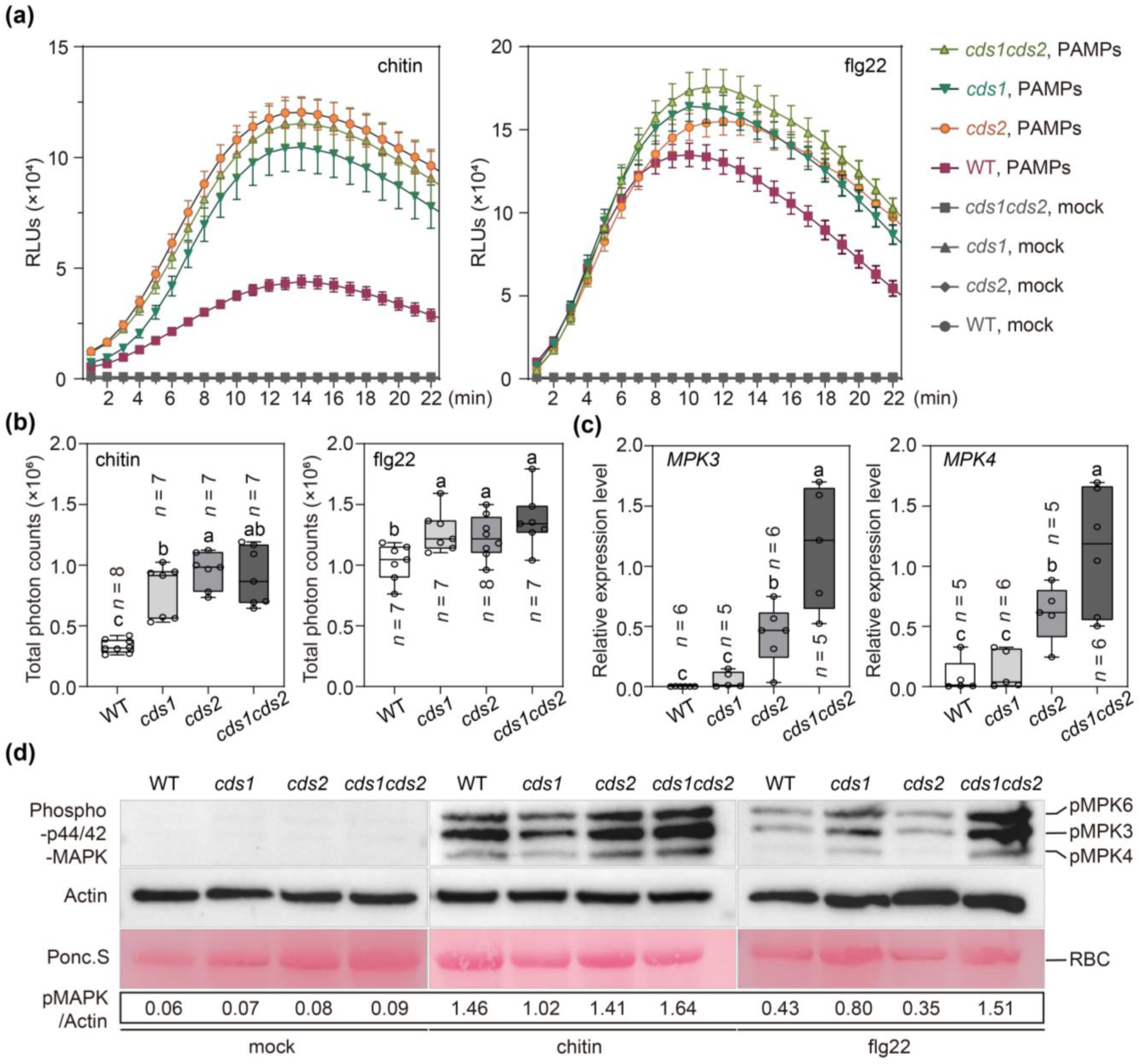
Plant immunity markers activated in *cds* mutants. (a) Reactive oxygen species (ROS) generation in *cds* mutant lines treated with chitin (120 nM) flg22 (200 nM) or water (mock); RLU, relative light unit. Leaves of 3-week-old *Arabidopsis* plants were used in the assays. Error bars represent mean ± SEM. (b) Total photon counts in *cds* mutant lines from (a). (c) RT-qPCR analyses of *MPK3* and *MPK4* expression levels in *cds* mutant lines. *ACTIN2* was used as the internal reference gene. (d) Immunoblot assays of MAPKs phosphorylation in *cds* leaves treated with chitin, flg22 and water (mock), respectively. RBC, ribulose-1,5-bis-phosphate carboxylase/oxygenase. Phosphorylated MAPKs were detected with the anti-phospho-p44/42 antibody, and goat anti-rabbit IgG HRP was used as the secondary antibody. Ponceau S staining shows the protein loading for each lane. Numbers below Ponceau S stain indicate the band intensity relative to that of Actin.

To investigate the underlying mechanisms by which *CDS* genes contribute to metabolism and defense, integrated transcriptomics, lipidomics, and metabolomics analyses of the *cds* mutant lines were conducted. To ensure the consistency of the results from the multi-omics association analysis, the leaves of *Arabidopsis* plants with the same growth condition were collected. Each analysis was performed with three biological replicates (Li et al., 2022). RNA-seq analysis showed that phospholipid biosynthesis and signaling genes, such as those involved in ‘fatty acid response’ and ‘lipid response’ were enriched in *cds1cds2* mutant lines. Signaling pathways associated with immunity, such as ‘fungal response’, ‘JA response’, and ‘response to oxidative stress’, were activated (Fig. 4a). The upregulated expression of these genes likely enhanced the resistance of the *cds1cds2* mutant to different pathogens. Further analysis revealed that a large number of genes associated with bacterial and fungal disease resistance responses were upregulated in the *cds1cds2* mutant lines. *WRKY* transcription factors, the important signaling factors for MAPK-mediated disease resistance responses, which regulate plant immune responses through promoting the expression of *SARD1* and *CBP60* in *Arabidopsis* (Chen et al., 2021), were upregulated in *cds1cds2.* BSK1 plays a key role in growth and immune regulation by binding to BRI1 (the receptor of brassinosteroid, which regulates plant growth and development) and FLS2 (the receptor that perceives flagellin 22 to activate a series of downstream immune responses) (Su et al., 2021). *GLYI4* is involved in the interaction between JA and SA pathways and enhances disease resistance in *Arabidopsis* (Proietti et al., 2019), and both genes were upregulated in *cds1cds2.* Our data suggest that *AtS40*, *NGAL2*, *RGL3*, and other genes associated with embryonic development and growth in plants were also affected in the *cds1cds2* mutant (Fig. 4b). *RGL3,* a negative regulator of gibberellic acid signaling that inhibits plant growth and development, was upregulated in *cds1cds2. AtS40* and *NGAL2* were also upregulated in *cds1cds2*; *AtS40* plays a conducive role in plant senescence, and *NGAL2* inhibits seed development (Fischer-Kilbienski et al., 2010, Wild and Achard, 2013, Zhang et al., 2015).

**FIGURE 4.**
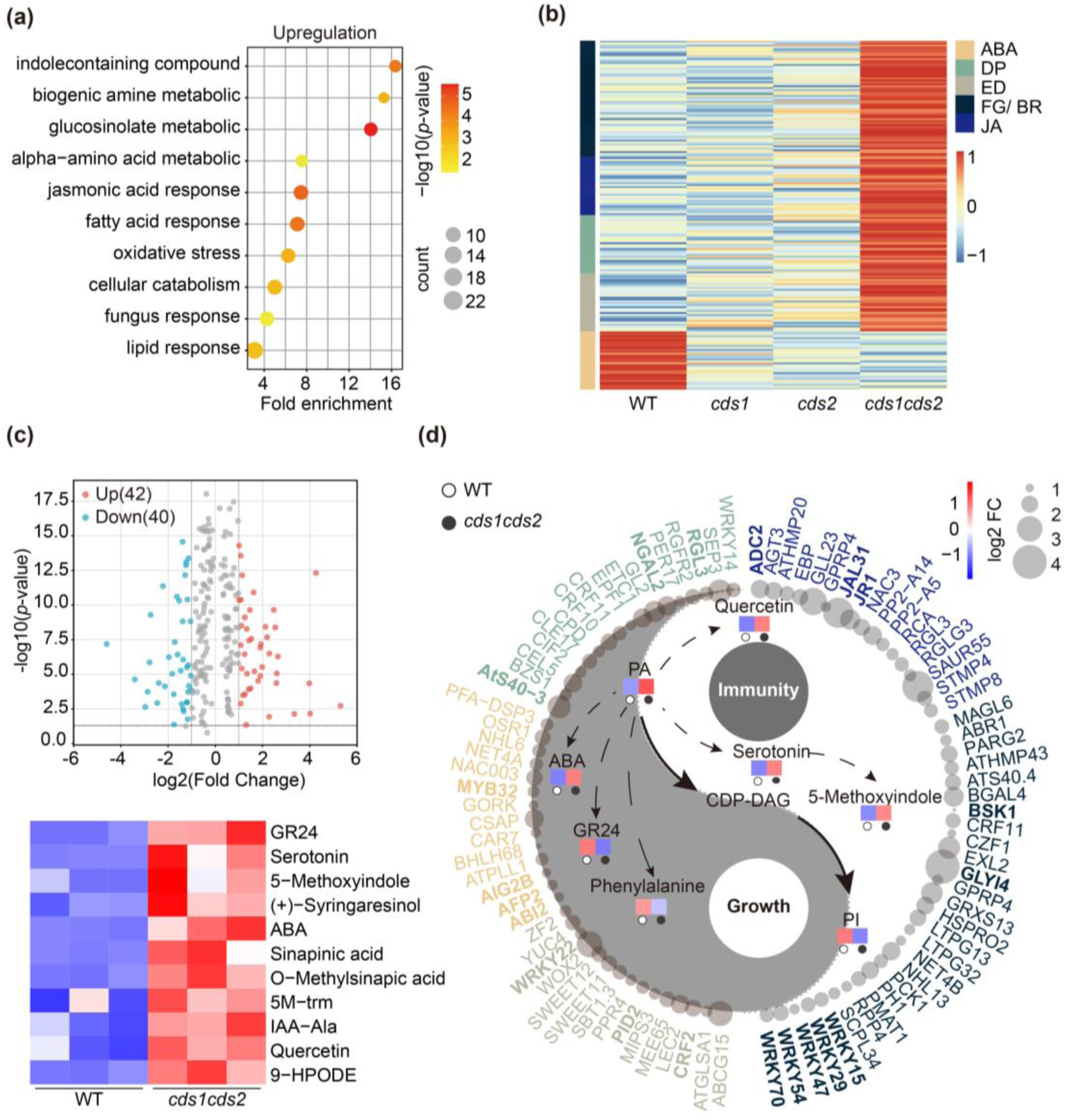
Integrated transcriptomics and metabolomics analyses of *cds1cds2*. (a) Gene ontology (GO) enrichment analysis of upregulated differentially expressed genes (DEGs) in *cds1cds2*. (b) Growth and immunity associated genes were upregulated in *cds1cds2*. ABA, abscisic acid response; DP, developmental process; ED, embryo development; FG/BR, fungus/ bacterium defense response; JA, jasmonic acid response. (c) Volcano plots of differential metabolites in *cds1cds2* vs. WT. (|Log2FC| ≥ 1.0, *p*-value < 0.05). The heat map shows the relative levels of metabolites associated with growth and immunity in WT and *cds1cds2* mutant lines. Each lane in heatmap shows the replicate of WT and *cds1cds2* mutant. (d) Differential metabolites and DEGs associated with plant growth and immunity in *cds1cds2*. The colors in the box indicate the relative amounts of metabolites in the WT and *cds1cds2* lines. The size of the circle indicates the relative expression levels of the gene.

Metabolomics analyses were conducted to further investigate the effects of *CDS* genes mutation on plant metabolism. These analyses revealed many changes in key metabolites that are likely involved in immunity and growth (Fig. 4c). The levels of serotonin, 5-Methoxyindole, and quercetin were elevated in the *cds1cds2* mutant (Fig. 4c). The accumulation of serotonin significantly enhances disease resistance in the rice *osspl5* mutant (Jin et al., 2015). 5-Methoxyindole, a congener of melatonin, inhibits the growth, formation, and conidial germination of the fungal pathogen *Fusarium graminearum* and exhibits potent anti-fungal activity (Kong et al., 2021). Quercetin, a plant hormone, suggests antimicrobial, antioxidant, and antiviral activities (Singh et al., 2021). Some metabolites with antibacterial activity, such as 9-HPODE, O-Methylsinapic acid, and sinapinic acid, were elevated in the *cds1cds2* mutants as potential plant phytoalexins (Chen, 2016, Zhao et al., 2019, Kimura and Yokota, 2004). The *cds1cds2* mutant showed a significant accumulation of ABA, an important phytohormone that plays a key physiological role in regulating seed development, growth inhibition, promoting flower, and fruit drop (Luo et al., 2014). Lipidomics results were consistent with previous reports that downregulation of *CDS1* and *CDS2* expression leads to PA accumulation and suppression of PI biosynthesis. As compared to WT, the relative content of PA was significantly higher, while the accumulation of PI was reduced in *cds1cds2* mutants. Both PA and PI have diverse roles, and altered levels of PA and PI might be a major cause of the alterations in the genes and metabolites mentioned above (Fig. 4d). The decreased levels of diacylglycerol phospholipids (DAG) and Lysophosphatidylglycero (LPG) were observed in *cds1cds2* mutant lines. The levels of Lysophosphatidylethanolamine (LPE), Lysophosphatidylcholine (LPC), Phosphatidylcholine (PC), as well as many other phospholipids, were increased in the *cds1cds2* mutant lines (Fig. S2). These transcriptomics and metabolomics results provide further evidence that *CDS* genes are negative regulators of immunity. Furthermore, the disruption of PA metabolic processes resulted in pleiotropic effects on growth and immunity, with a wide range of transcriptomics and metabolic changes.

## 4. DISCUSSION

Previous studies have shown that loss of function of one of the *AtCDS1* and *AtCDS2* genes does not significantly affect plant growth. Complete knockout of both *AtCDS1* and *AtCDS2* genes results in seedling lethality in *Arabidopsis* (Zhou et al., 2013), suggesting these genes are functionally redundant. Recently, *Arabidopsis* T-DNA insertional mutants with partial loss of *CDS1* and *CDS2* function are characterized, exhibiting defects in embryonic development (Du et al., 2022). Here, we showed that the *cds1cds2* double mutant exhibited significantly impaired growth, manifesting in reduced plant height and diminished yield compared to the WT (Fig. 1). These phenotypic changes may be attributed to defects in early embryonic development. Our results suggested that the *cds1cds2* mutant exhibited enhanced in broad-spectrum resistance (Fig. 2) and displayed more robust immune response compared to the WT (Fig. 3). These findings were consistent with previous results demonstrating that the *cds1cds2* mutant shows enhanced resistance to oomycetes. In rice, mutation of the CDP-DAG synthase gene *RBL1* results in lesion mimic phenotype and confers broad-spectrum disease resistance (Sha et al., 2023). Similarly, mutations of the rice *OsCDS5* gene enhances ROS production and promotes expression levels of multiple defense-related genes, and the *oscds5* mutant exhibits enhanced resistance to rice blast, bacterial blight, and bacterial leaf streak (Sun et al., 2024). Through our studies in rice and *Arabidopsis*, we implied that partial loss of function of the *CDS* genes enhances plant resistance to diverse pathogens, underscoring the conserved roles of CDP-DAG synthase in regulating plant immunity.

The enhanced immunity of the *cds1cds2* mutant could be a result of metabolic changes in multiple pathways. Previous findings indicate that PA, a signaling molecule in plant immunity (Jennings and Epand, 2020), interacts with RBOH, subsequently activates downstream MPK3 and MPK6, promoting ROS production and MAPK phosphorylation to enhance disease resistance (Kong et al., 2024). The level of PA was increased, as was the production of ROS and the levels of MAPK phosphorylation in the *cds1cds2* mutant, suggested the partial loss of CDS1 and CDS2 function in the *cds1cds2* mutant led to the accumulation of PA, which activated the downstream ROS and MAPK pathways and conferred disease resistance in the *cds1cds2* mutant. Alternatively, the enhanced immunity observed in the *cds1cds2* mutant may be attributed to the reduced PI levels in *Arabidopsis.* PI derivatives are modulated by plant pathogens to facilitate infection and are considered disease susceptibility factors. One of the PI derivates, PI(4,5)P_2_, is recruited to the extrahaustorial membrane at powdery mildew infection sites to facilitate infection (Qin et al., 2020). Mutation of PI(4,5)P_2_ biosynthetic enzymes causes reduced PI(4,5)P_2_ levels and enhanced resistance. In the *rbl1* rice mutant, reduced levels of PI and PIPs are observed, and the formation of cellular structures involved in pathogen effector protein translocation is disrupted, thereby resulting in enhanced immunity to multiple pathogens (Sha and Li, 2023). The level of PIPs requires to be measured, and the role of PIPs in enhanced immunity of the *cds1cds2* mutant requires further investigation. LPE, a natural occurring phospholipid, has been demonstrated to play a significant role in modulating early stages of plant senescence, primarily by delaying senescence-associated processes. LPE-treatment induces the signaling of SA and diverse immune defense responses, enhancing resistance to *Pst* DC3000 of *Arabidopsis* (Völz et al., 2021). The increased levels of LPE and SA in the *cds1cds2* mutants suggested that the enhanced resistance in the *cds1cds2* mutants was achieved by accumulating LPE to activate the SA signaling pathway (Fig. S1-S2). Finally, the enhanced resistance observed in *Arabidopsis cds1cds2* mutant is potentially linked to the upregulation of immunity-related genes and the increased accumulation of defense-associated metabolites. Our integrated transcriptomics, lipidomics, and metabolomics analyses showed that the *WRKY* transcription factors, *BSK1,* and *GLYI4* were significantly upregulated in *cds1cds2* plants, which reportedly activated JA and SA signaling pathways and enhanced disease resistance in *Arabidopsis* (Fig. 4b, d). The significant accumulation of SA in *cds1cds2* mutants corroborated the activation of SA-mediated immune responses, underscoring the role of SA as a key signaling molecule (Fig. S1). Serotonin and 5-Methoxyindole were elevated in *cds1cds2,* which are plant defense-associated metabolites (Fig. 4c, d). Our findings suggest that the enhanced immunity observed in *Arabidopsis cds1cds2* mutant was attributable to multiple factors including elevated ROS production, increased MAPK phosphorylation, reduced PI levels, and upregulation of defense-related genes and metabolites.

Knock-down mutations of both *Arabidopsis CDS1* and *CDS2* genes enhanced plant immunity but reduced plant growth. In rice, the partially functional *CDS* gene confers broad-spectrum disease resistance without compromising yield (Sha et al., 2023). Additionally, by editing the upstream open reading frame (uORF) of the gene, it may be possible to regulate protein abundance. For example, the introduction of uORF regulatory elements with inhibitory activities into the immunoregulatory factor of *Arabidopsis* modulates the protein translation level of AtNPR1, thereby improving broad-spectrum plant disease resistance without affecting growth and development (Zhang et al., 2018). Using such strategies, we anticipate that fine-tuning *CDS* transcript or protein abundance to balance growth and immunity in crops could then be applied to breeding programs for crops as novel sources of disease resistance.

## ACKNOWLEDGMENTS

We thank Prof. Long Yang at Huazhong Agricultural University for the *B. cinerea* strain. We thank Prof. Xiufang Xin at CAS Center for Excellence in Molecular Plant Sciences for the *Pst* DC3000 bacterial strains. We thank Dr. Ricky J. Milne at CSIRO, Australia for critical reading of this manuscript. We thank Anum Bashir and Xinyu Han for their help in revising this manuscript. This work was supported by STI2030-Major Projects (2023ZD04070), the Key R&D Program of Hubei Province (2023BBB171), National Natural Science Foundation of China (32172373), Fundamental Research Funds for the Central Universities (2662023PY006 and AML2023A05), and the Open Research Fund of the State Key Laboratory of Hybrid Rice (Wuhan University) (KF202202) to G.L. G.S. is supported by the Open Research Fund of the Guangdong Province Key Laboratory of Microbial Signals and Disease Control (MSDC2023-02). This work was also supported by Hubei Hongshan Laboratory.

## AUTHOR CONTRIBUTIONS

G.L. and G.S. designed the experiments. J.C. provided the *S. sclerotiorum* strain. R.T., X.L. and J.C. performed pathogen infection assays on *Arabidopsis*. G.S., Q.G. and R.T. performed MAPK phosphorylation assays. H.X. and X.D, generated transgenic plants. R.T. performed *Arabidopsis* growth phenotypic analyses, ROS and RT-qPCR assays. R.T., Q.G. and Q.L. performed lipid extraction and lipidomics analysis. G.S., W.Y., L.Y. and R.T. performed bioinformatics and RNA-seq data analyses. G.S., R.T., Y.L. and J.L. performed metabolomics profiling analyses and SA analyses. G.L., G.S. and R.T. drafted the manuscript. G.L., G.S., Q.G. and R.T. revised the manuscript with inputs from all authors. All authors read and approved the final manuscript.

## DATA AVAILABILITY

The genes mentioned in this study are as follows: *AtCDS1* (AT1G62430, CP138171.1); *AtCDS2* (AT4G22340, CP138174.1). RNA-seq data are available in the National Center for Biotechnology Information (NCBI) (http://www.ncbi.nlm.nih.gov/) under the accession number of PRJNA1091892.

## SUPPORTING INFORMATION

Additional Supporting Information may be found at the end of the article.

### SUPPLEMENTAL FIGURES

**Figure S1.**
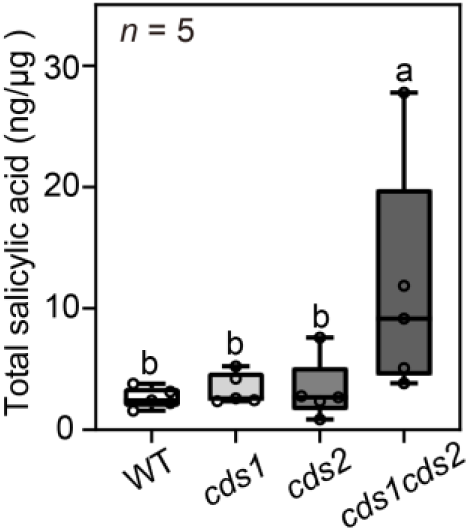
Total salicylic acid levels of wild-type and *cds* mutants. Salicylic acid levels were significantly increased in the *cds1cds2* mutant. Total SA was isolated from leaves of 4-week-old *Arabidopsis* seedlings. Data are displayed as box and whisker plots with individual data points. The box plot elements are: center line, median; box limits, 25^th^ and 75^th^ percentiles. *n* indicates the number of biological replicates. Letters indicate statistically significant differences (ANOVA, *p*< 0.05). Duncan’s Multiple Range Test as method of post-hoc test.

**Figure S2.**
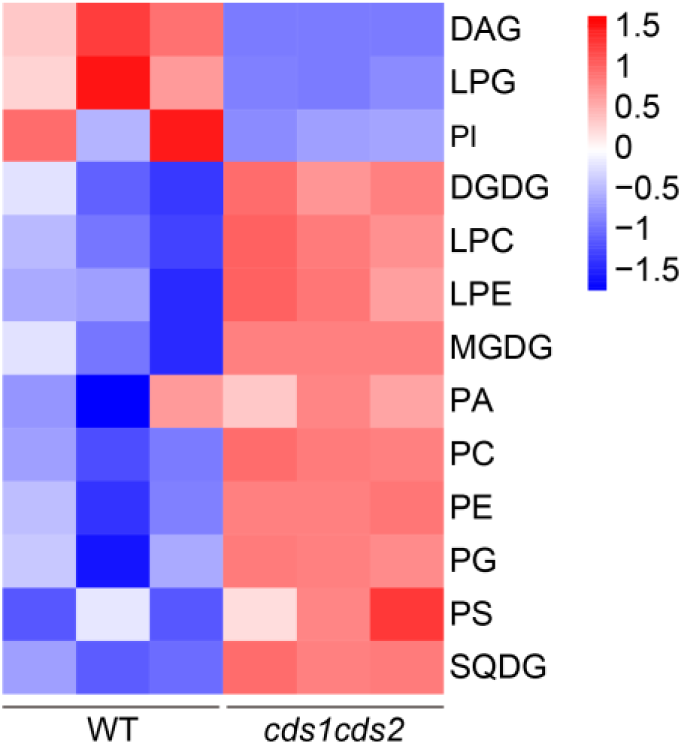
Lipidomics analyses of WT and *cds1cds2* mutant lines. The heat map shows the lipids composition and molecular species composition of WT and *cds1cds2* mutant lines. Lipids were extracted and analyzed from leaves of 4-week-old *Arabidopsis* seedlings. Labeling of the corresponding lipids from WT and *cds1cds2* mutant lines in per cent of total labeling. Each lane in heatmap shows the replicate of WT and *cds1cds2* mutant. DAG, Diacylglycerol Phospholipids; LPG, Lysophosphatidylglycerol; PI, phosphatidylinositol; DGDG, digalactosyldiacylglycerol; LPC, Lysophosphatidylcholine; LPE, Lyso-phosphatidylethanolamine; MGDG, monogalactosyldiacylglycerol; PA, phosphatidic acid; PC, phosphatidylcholine; PE, phosphatidylethanolamine; PG, phosphatidylglycerol; PS, Phosphatidylserine; SQDG, sulfoquinovosyldiacylglycerol.

**Supplementary Table S1.**
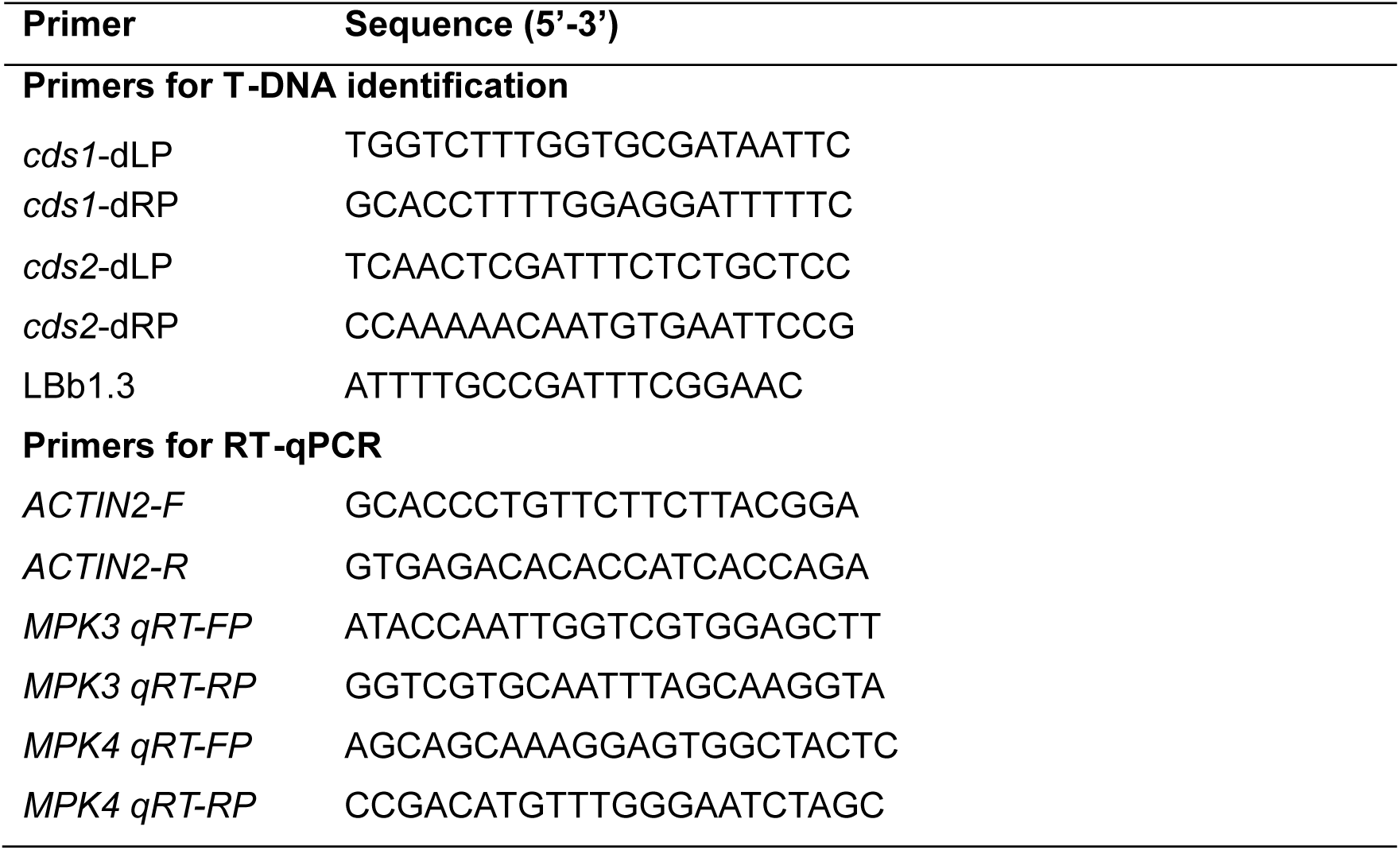
Primers used in this study.

## References

Ali, U., Lu, S., Fadlalla, T., Iqbal, S., Yue, H., Yang, B., et al. (2022) The functions of phospholipases and their hydrolysis products in plant growth, development and stress responses. Prog Lipid Res. 86, 101158.

Chen, C. (2016) Sinapic acid and its derivatives as medicine in oxidative stress-induced diseases and aging. Oxid Med Cell Longev. 2016, 3571614.

Chen, S., Ding, Y., Tian, H., Wang, S. & Zhang, Y. (2021) WRKY54 and WRKY70 positively regulate *SARD1* and *CBP60g* expression in plant immunity. Plant Signal Behav. 16, 1932142.

Du, X. Q., Yao, H. Y., Luo, P., Tang, X. C. & Xue, H. W. (2022) Cytidinediphosphate diacylglycerol synthase-mediated phosphatidic acid metabolism is crucial for early embryonic development of *Arabidopsis*. PLoS Genet. 18, e1010320.

Fischer-Kilbienski, I., Miao, Y., Roitsch, T., Zschiesche, W., Humbeck, K. & Krupinska, K. (2010) Nuclear targeted AtS40 modulates senescence associated gene expression in *Arabidopsis thaliana* during natural development and in darkness. Plant Mol Biol. 73, 379–90.

Gong, Y., Fu, Y., Xie, J., Li, B., Chen, T., Lin, Y., et al. (2022) *Sclerotinia sclerotiorum SsCut1* modulates virulence and cutinase activity. J Fungi 8.

Haselier, A., Akbari, H., Weth, A., Baumgartner, W. & Frentzen, M. (2010) Two closely related genes of *Arabidopsis* encode plastidial cytidinediphosphate diacylglycerol synthases essential for photoautotrophic growth. Plant Physiol. 153, 1372–84.

Hong, Y., Yuan, S., Sun, L., Wang, X. & Hong, Y. (2018) Cytidinediphosphate-diacylglycerol synthase 5 is required for phospholipid homeostasis and is negatively involved in hyperosmotic stress tolerance. Plant J. 94, 1038–1050.

Hu, Y., Ding, Y., Cai, B., Qin, X., Wu, J., Yuan, M., et al. (2022) Bacterial effectors manipulate plant abscisic acid signaling for creation of an aqueous apoplast. Cell Host Microbe. 30, 518–529.e6.

Huang, J., Liang, X., Xuan, Y., Geng, C., Li, Y., Lu, H., et al. (2017) A reference human genome dataset of the BGISEQ-500 sequencer. Gigascience. 6, 1–9.

Jennings, W. & Epand, R. M. (2020) CDP-diacylglycerol, a critical intermediate in lipid metabolism. Chem Phys Lipids. 230, 104914.

Jin, B., Zhou, X., Jiang, B., Gu, Z., Zhang, P., Qian, Q., et al. (2015) Transcriptome profiling of the *spl5* mutant reveals that *SPL5* has a negative role in the biosynthesis of serotonin for rice disease resistance. Rice. 8, 18.

Kamaruzzaman, M., He, G., Wu, M., Zhang, J., Yang, L., Chen, W., et al. (2019) A novel partitivirus in the hypovirulent isolate QT5-19 of the plant pathogenic fungus *Botrytis cinerea*. Viruses. 11.

Kimura, H. & Yokota, K. (2004) Characterization of metabolic pathway of linoleic acid 9-hydroperoxide in cytosolic fraction of potato tubers and identification of reaction products. Appl Biochem Biotechnol. 118, 115–32.

Kong, L., Ma, X., Zhang, C., Kim, S. I., Li, B., Xie, Y., et al. (2024) Dual phosphorylation of DGK5-mediated PA burst regulates ROS in plant immunity. Cell. 187, 609–623.e21.

Kong, M., Liang, J., Ali, Q., Wen, W., Wu, H., Gao, X., et al. (2021) 5-Methoxyindole, a chemical homolog of melatonin, adversely affects the phytopathogenic fungus *Fusarium graminearum*. International Journal of Molecular Sciences. 22, 10991.

Li, J. & Wang, X. (2019) Phospholipase D and phosphatidic acid in plant immunity. Plant Sci. 279, 45–50.

Li, J., Yao, S., Kim, S. C. & Wang, X. (2024) Lipid phosphorylation by a diacylglycerol kinase suppresses ABA biosynthesis to regulate plant stress responses. Mol Plant. 17, 342–358.

Li, Y., Yang, Z., Yang, C., Liu, Z., Shen, S., Zhan, C., et al. (2022) The *NET* locus determines the food taste, cooking and nutrition quality of rice. Sci Bull. 67, 2045–2049.

Liu, Q., Ning, Y., Zhang, Y., Yu, N., Zhao, C., Zhan, X., et al. (2017) OsCUL3a negatively regulates cell death and immunity by degrading OsNPR1 in rice. Plant Cell. 29, 345–359.

Livak, K. J. & Schmittgen, T. D. (2001) Analysis of relative gene expression data using real-time quantitative PCR and the 2 -^ΔΔcT^ method. Methods. 25, 402–8.

Lu, S., Liu, H., Jin, C., Li, Q. & Guo, L. (2019) An efficient and comprehensive plant glycerolipids analysis approach based on high-performance liquid chromatography-quadrupole time-of-flight mass spectrometer. Plant Direct. 3, e00183.

Luo, X., Chen, Z., Gao, J. & Gong, Z. (2014) Abscisic acid inhibits root growth in *Arabidopsis* through ethylene biosynthesis. Plant J. 79, 44–55.

Proietti, S., Falconieri, G. S., Bertini, L., Baccelli, I., Paccosi, E., Belardo, A., et al. (2019) GLYI4 plays a role in methylglyoxal detoxification and jasmonate-mediated stress responses in *Arabidopsis thaliana*. Biomolecules. 9, 635.

Qi, F., Li, J., Ai, Y., Shangguan, K., Li, P., Lin, F., et al. (2024) DGK5β-derived phosphatidic acid regulates ROS production in plant immunity by stabilizing NADPH oxidase. Cell Host Microbe. 32, 425–440.e7.

Qin, L., Zhou, Z., Li, Q., Zhai, C., Liu, L., Quilichini, T. D., et al. (2020) Specific recruitment of phosphoinositide species to the plant-pathogen interfacial membrane underlies *Arabidopsis* susceptibility to fungal infection. Plant Cell 32, 1665–1688.

Sha, G. & Li, G. (2023) Effector translocation and rational design of disease resistance. Trends Microbiol. 31, 1202–1205.

Sha, G., Sun, P., Kong, X., Han, X., Sun, Q., Fouillen, L., et al. (2023) Genome editing of a rice CDP-DAG synthase confers multipathogen resistance. Nature. 618, 1017–1023.

Singh, P., Arif, Y., Bajguz, A. & Hayat, S. (2021) The role of quercetin in plants. Plant Physiol Biochem. 166, 10–19.

Su, B., Zhang, X., Li, L., Abbas, S., Yu, M., Cui, Y., et al. (2021) Dynamic spatial reorganization of BSK1 complexes in the plasma membrane underpins signal-specific activation for growth and immunity. Mol Plant. 14, 588–603.

Sun, Q., Xiao, Y., Song, L., Yang, L., Wang, Y., Yang, W., et al. (2024) Mutation of *OsCDS5* confers broad-spectrum disease resistance in rice. Molecular Plant Pathology. 25, e13430.

Völz, R., Park, J. Y., Harris, W., Hwang, S. & Lee, Y. H. (2021) Lyso-phosphatidylethanolamine primes the plant immune system and promotes basal resistance against hemibiotrophic pathogens. BMC Biotechnol. 21, 12.

Wild, M. & Achard, P. (2013) The DELLA protein RGL3 positively contributes to jasmonate/ethylene defense responses. Plant Signal Behav. 8, e23891.

Xing, J., Zhang, L., Duan, Z. & Lin, J. (2021) Coordination of phospholipid-based signaling and membrane trafficking in plant immunity. Trends Plant Sci. 26, 407–420.

Yao, S., Kim, S. C., Li, J., Tang, S. & Wang, X. (2024) Phosphatidic acid signaling and function in nuclei. Prog Lipid Res. 93, 101267.

Yu, G., Wang, L. G., Han, Y. & He, Q. Y. (2012) clusterProfiler: an R package for comparing biological themes among gene clusters. OMICS. 16, 284–7.

Yuan, M., Jiang, Z., Bi, G., Nomura, K., Liu, M., Wang, Y., et al. (2021) Pattern-recognition receptors are required for NLR-mediated plant immunity. Nature. 592, 105–109.

Zhai, K., Liang, D., Li, H., Jiao, F., Yan, B., Liu, J., et al. (2022) NLRs guard metabolism to coordinate pattern- and effector-triggered immunity. Nature. 601, 245–251.

Zhang, F., Guo, H., Huang, J., Yang, C., Li, Y., Wang, X., et al. (2020) A UV-B-responsive glycosyltransferase, Osugt706c2, modulates flavonoid metabolism in rice. Sci China Life Sci. 63, 1037–1052.

Zhang, H., Si, X., Ji, X., Fan, R., Liu, J., Chen, K., et al. (2018) Genome editing of upstream open reading frames enables translational control in plants. Nature Biotechnology. 36, 894–898.

Zhang, Y., Du, L., Xu, R., Cui, R., Hao, J., Sun, C., et al. (2015) Transcription factors SOD7/NGAL2 and DPA4/NGAL3 act redundantly to regulate seed size by directly repressing *KLU* expression in *Arabidopsis thaliana*. Plant Cell 27, 620–32.

Zhang, Y., Zhu, H., Zhang, Q., Li, M., Yan, M., Wang, R., et al. (2009) Phospholipase Dα1 and phosphatidic acid regulate NADPH oxidase activity and production of reactive oxygen species in ABA-mediated stomatal closure in *Arabidopsis*. Plant Cell. 21, 2357–77.

Zhao, J., Chen, Y., Ding, Z., Zhou, Y., Bi, R., Qin, Z., et al. (2024) Identification of propranolol and derivatives that are chemical inhibitors of phosphatidate phosphatase as potential broad-spectrum fungicides. Plant Commun. 5, 100679.

Zhao, Z., Song, H., Xie, J., Liu, T., Zhao, X., Chen, X., et al. (2019) Research progress in the biological activities of 3,4,5-trimethoxycinnamic acid (TMCA) derivatives. European Journal of Medicinal Chemistry. 173, 213–227.

Zhou, J., Liu, C., Chen, Q., Liu, L., Niu, S., Chen, R., et al. (2022) Integration of rhythmic metabolome and transcriptome provides insights into the transmission of rhythmic fluctuations and temporal diversity of metabolism in rice. Sci China Life Sci. 65, 1794–1810.

Zhou, Y., Peisker, H., Weth, A., Baumgartner, W., Dörmann, P. & Frentzen, M. (2013) Extraplastidial cytidinediphosphate diacylglycerol synthase activity is required for vegetative development in *Arabidopsis thaliana*. Plant J. 75, 867–79.

